# The ubiquitylome of developing cortical neurons

**DOI:** 10.1101/2020.10.13.337782

**Authors:** Shalini Menon, Dennis Goldfarb, Emily M. Cousins, M. Ben Major, Stephanie L. Gupton

## Abstract

TRIM9 and TRIM67 are neuronally-enriched E3 ubiquitin. Both genes are required for neuronal morphological responses to the axon guidance cue netrin-1. For example, our previously published work demonstrated that the actin polymerase VASP and the netrin receptor DCC exhibit TRIM9 dependent ubiquitylation that is lost upon netrin stimulation. Deletion of either gene in the mouse results in subtle neuroanatomical anomalies yet overt deficits in spatial learning and memory. Despite their role in neuronal form and function, the identity of few TRIM9 or TRIM67 substrates are known. Here we performed ubiquitin remnant profiling approach in cultured wildtype and knockout murine embryonic cortical neurons to identify ubiquitylated peptides and proteins, with the ultimate goal of identifying substrates of TRIM9 and TRIM67 that exhibited reduced ubiquitylation in the absence of the ligase. This work reveals the ubiquitylome of developing cortical neurons.

## Introduction

Ubiquitylation is a posttranslational protein modification, in which the small protein ubiquitin is covalently attached to substrate proteins. This modification can mark substrates for degradation by the proteasome or lysosomes, or alternatively can alter protein localization, function, trafficking, and protein-protein interactions. Ubiquitylation is a ubiquitous modification, but it is enriched in the brain and often accumulates in neurodegenerative conditions (Fischer et al., 2009).

TRIM9 and TRIM67 are neuronally-enriched E3 ubiquitin ligases (Boyer et al., 2018a; Berti et al., 2002). Both genes are required for neuronal morphological responses to the axon guidance cue netrin-1 (Winkle et al., 2014; Boyer et al., 2018; Plooster et al., 2017; Boyer et al., 2020; Menon et al., 2015; Urbina et al., 2018; Winkle et al., 2016). For example, our previously published work demonstrated that the actin polymerase VASP and the netrin receptor DCC exhibit TRIM9 dependent ubiquitylation that is lost upon netrin stimulation (Menon et al., 2015; Plooster et al., 2017; Boyer et al., 2020). Deletion of either gene in the mouse results in subtle neuroanatomical anomalies yet overt deficits in spatial learning and memory (Menon et al., 2015; Winkle et al., 2016; Boyer et al., 2018). Despite their role in neuronal form and function, the identity of few TRIM9 or TRIM67 substrates are known. Here we performed ubiquitin remnant profiling approach (Xu et al., 2010) in cultured wildtype and knockout murine embryonic cortical neurons to identify ubiquitylated peptides and proteins, with the ultimate goal of identifying substrates of TRIM9 and TRIM67 that exhibited reduced ubiquitylation in the absence of the ligase. To identify substrates that show either netrin-dependent or netrin-sensitive ubiquitination, experiments were done with or without supplementation of cultures with the guidance cue netrin-1.

## Results

Ubiquitylated peptides were enriched from cultured cortical neurons from *Trim9^+/+^/Trim67^+/+^, Trim9^-/-^/Trim67^+/+^, Trim9^+/+^/Trim67^-/-^, Trim9^-/-^/Trim67^-/-^* embryos at 2.5 days in vitro (DIV) with or without 40 minutes of netrin-1 supplementation (**Figure 1A)**. The enriched peptides were analyzed by LC/MS/MS. This generated an extensive list of 854 unique proteins and 1454 unique peptides across all genotypes and treatments that were ubiquitylated during early neuronal development (Table 1, sheet 1: list of ubiquitylated proteins, sheet 2: all unique peptides with modified lysine). Although no differences in ubiquitylation between any condition reached statistical significance, likely due to batch variability between replicates and insufficient sample size, these results revealed the vast array of protein ubiquitylation during this developmental window.

**Figure 1.**
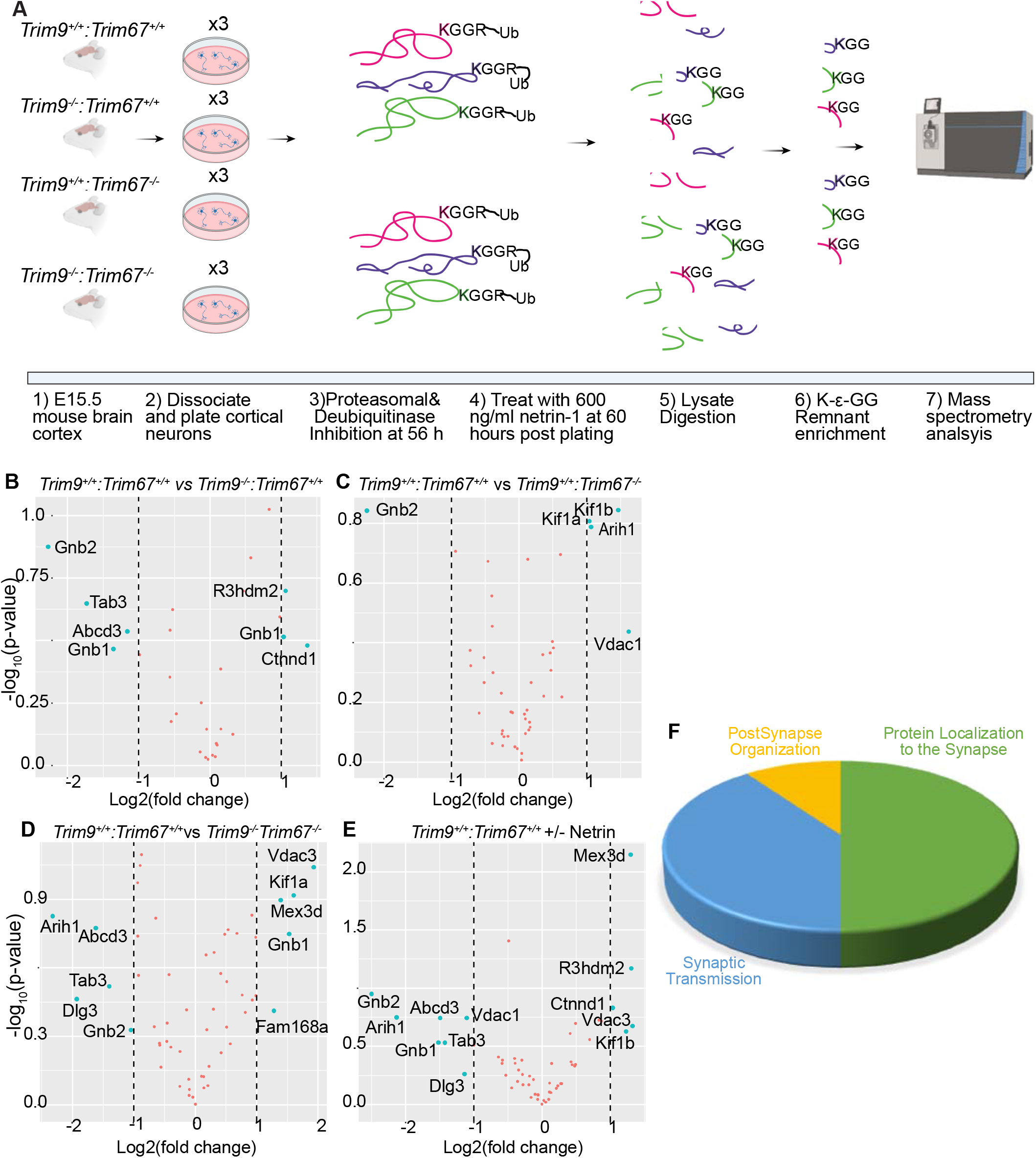
The ubiquitylome of developing cortical neurons. **A)** Graphical representation of approach. Embryonic day 15.5 (E15.5) neurons were dissected from the cortex of mice with indicated genotypes, and cultured for 56 hours prior to addition of proteasomal and deubiquitinase inhibitors for 4 hrs. Half of the plated neurons were treated with 600ng/ml of netrin-1 for 40 min prior to lysis. Neurons were lysed and trypsin digested. Ubiquitylated peptides were enriched and analyzed by mass spectrometry. **B-E)** Volcano plots demonstrating changes in ubiquitylation of proteins that overlapped with high-confidence TRIM9 and TRIM67 interaction partners between the indicated conditions. **F)** Pie-chart of biological processes enriched via Gene Ontology (GO) classification of proteins that overlapped between high-confidence TRIM9 and TRIM67 interaction partners and altered DiGly enrichment.

In a previous study, we identified a putative TRIM9 and TRIM67 interactome using proximity-dependent labeling with the promiscuous biotin ligase (BirA*) attached to either TRIM9 or TRIM67 (Menon et al., 2020). A comparison of the ubiquitylated proteome from developing brains with the top 0.25% of the interaction candidates identified peptides from 55 proteins that showed an increase or decrease (albeit insignificant) in ubiquitylation levels in the absence of *Trim9, Trim67*, or both relative to wildtype (Figure 1B-D). Similar analysis revealed proteins with altered ubiquitylation upon netrin application. Gene ontology analysis of these 55 proteins demonstrated synaptic enrichment, with only three categories identified: protein localization to the synapse, synaptic transmission, and postsynapse organization. This is remarkable as both proteomic approaches were done in cortical neurons well before synapses formed. However a synaptic function for TRIM9 and TRIM67 is consistent with behavioral phenotypes identified in knockout animals(Winkle et al., 2016; Boyer et al., 2018). 8 and 5 proteins were identified as candidate TRIM9 and TRIM67 substrates, respectively. Proteins that exhibited differential ubiquitylation upon loss of *Trim9* included Adcd3, Ctnnd1, Gnb1, Gnb2, R3hdm2, Tab3. Proteins that exhibited differential ubiquitylation upon loss of *Trim67* included Arih1, Gnb2, Kif1a, Kif1b, Vdac1. Similarly, Abcd3, Arih1, Dlg3, Gnb1, Gnb2, Kif1a, Mex3d, Tab3, Tcrp1, Vdac3 were found to be differentially ubiquitylated in *Trim9^-/-^* and *Trim67^-/-^* neurons relative to *Trim9^+/+^:Trim67^+/+^* neurons. Further exploration will be needed to validate TRIM9 and TRIM67 dependent ubiquitylation of these proteins during neuronal development. The identification of ubiquitylated peptides and proteins in developing cortical neurons will be illuminating for many future studies.

## Methods

### Animals

All mouse lines were on a C57BL/6J background and bred at UNC with approval from the Institutional Animal Care and Use Committee. Timed pregnant females were obtained by placing male and female mice together overnight; the following day was designated as E0.5 if the female had a vaginal plug. Generation of *Trim9^-/-^* and *Trim67^-/-^* mice has been described previously (Boyer et al., 2018; Winkle et al., 2014).

### Cortical neuron culture

E15.5 litters were removed from wildtype, *Trim9^-/-^, Trim67^-/-^* and *Trim9^-/-^:Trim67^-/-^* pregnant dams by postmortem cesarean section. Dissociated cortical neuron cultures were prepared as described previously (Kwiatkowski et al., 2007). Briefly, cortices were micro-dissected, cortical neurons were dissociated with 0.25% trypsin for 20 min at 37°C, followed by quenching of trypsin with Trypsin Quenching Medium (TQM, neurobasal medium supplemented with 10% FBS and 0.5 mM glutamine). After quenching, cortices were gently triturated in neurobasal medium supplemented with TQM. Dissociated neurons were then pelleted at 100 g for 7 min. The pelleted cells were gently resuspended in Serum Free Medium (SFM, neurobasal medium supplemented with B27 (Invitrogen) and plated on tissue culture plastic coated with 1 mg/ml poly-D-lysine (Sigma-Aldrich).

### Enrichment of ubiquintated peptides

Primary cortical neurons from *Trim9^+/+^/Trim67^+/+^* (wildtype,WT), *Trim9^-/-^/Trim67^+/+^* (*Trim9* null), *Trim9^+/+^/Trim67^-/-^* (*Trim67* null), *Trim9^-/-^/Trim67^-/-^* (*Trim9/Trim67* double knockout, 2KO) murine E15.5 litters were dissociated and cultured on Poly-D-Lysine (Sigma) treated dishes. Approximately 56 hrs post-plating, cells were treated with 100 nM Bortezomib and 5 μM PR-619 for 4 hrs. 40 min prior to collecting cells, half of all plated cells were treated with 600 ng/ml of Netrin. Cells were harvested and stored at −80°C until dissociated neurons from 20-25 pups per genotype per condition were collected. The harvested cells were lysed for 30 min on ice in 1000 μl of freshly prepared buffer (8 M urea in 50 mM Tris-HCl pH7.5, 150 mM NaCl, 1 mM EDTA, 2 μg/ml apoprotinin, 10 μg/ml leupeptin, 1 mM PMSF, 50 μM PR-619 and 1 mM Iodoacetamide). Lysates were sonicated 3x 30 s cycle at 10% output, and clarified at 21.1k g at 4°C for 15 min. Clarified lysates were then reduced with 5 mM DTT at room temperature for 45 min followed by alkylation for 30 min at room temperature using 10 mM chloroacetamide. The reduced and alkylated protein lysates were diluted using 50 mM Tris-Cl to obtain a final concentration of 2 M urea in solution followed by overnight digestion with trypsin (1:1000)(Promega). Peptides were then desalted using c-18 Sep-Pack columns (Waters). Eluted peptides are lyophilized following which immunoaffinity purification was performed using the PTMScan Ubiquitin Remnant Motif (K-GG) Kit. The affinity purified peptides were desalted using C18 columns (Pierce).

### Mass spectrometry and data analysis

Reverse-phase nano-high-performance liquid chromatography (nano-HPLC) coupled with a nanoACQUITY ultraperformance liquid chromatography (UPLC) system (Waters Corporation; Milford, MA) was used to separate trypsinized peptides. Trapping and separation of peptides were performed in a 2 cm column (Pepmap 100; 3-m particle size and 100-Å pore size), and a 25-cm EASYspray analytical column (75-m inside diameter [i.d.], 2.0-m C18 particle size, and 100-Å pore size) at 300 nL/min and 35°C, respectively. Analysis of a 150-min. gradient of 2% to 25% buffer B (0.1% formic acid in acetonitrile) was performed on an Orbitrap Fusion Lumos mass spectrometer (Thermo Scientific). The ion source was operated at 2.4kV and the ion transfer tube was set to 300°C. Full MS scans (350-2000 m/z) were analyzed in the Orbitrap at a resolution of 120,000 and 1e6 AGC target. The MS2 spectra were collected using a 1.6 m/z isolation width and were analyzed either by the Orbitrap or the linear ion trap depending on peak charge and intensity using a 3 s TopSpeed CHOPIN method. Orbitrap MS2 scans were acquired at 7500 resolution, with a 5e4 AGC, and 22 ms maximum injection time after HCD fragmentation with a normalized energy of 30%. Rapid linear ion trap MS2 scans were acquired using an 4e3 AGC, 250 ms maximum injection time after CID 30 fragmentation. Precursor ions were chosen based on intensity thresholds (>1e3) from the full scan as well as on charge states (2-7) with a 30-s dynamic exclusion window. Polysiloxane 371.10124 was used as the lock mass.

For the data analysis all raw mass spectrometry data were searched using MaxQuant version 1.5.7.4. Search parameters were as follows: UniProtKB/Swiss-Prot mouse canonical sequence database (downloaded 1 Feb 2017), static carbamidomethyl cysteine modification, specific trypsin digestion with up to two missed cleavages, variable protein N-terminal acetylation and methionine oxidation, digly variable modification (+114.043), match between runs, and label-free quantification (LFQ) with a minimum ratio count of 2. Digly peptide intensities were log2 transformed and median normalized between experiments. The mass spectrometry proteomics data have been deposited to the ProteomeXchange Consortium via the PRIDE (Perez-Riverol et al., 2019) partner repository with the dataset identifier PXD021818.

### Data representation and Gene Ontology (GO) analysis

Gene ontology enrichment analysis of the neuronal ubiquitinome was performed using the ClueGo application available for download in the Cytoscape_3.7.

## Author Contributions

SM: Project design; data acquisition, curation, and validation, data analysis, data interpretation; training undergraduate researchers, manuscript and figure preparation; DG: Mass spectrometry (MS) and MS data analysis; EMC: Sample preparation and MS MBM: Oversaw MS methods and MS data analysis; manuscript editing; SLG: Project conceptualization, design, administration, funding acquisition; interpretation of data; manuscript and figure preparation and approval.

## Acknowledgements

We thank Vong Thoong, Caroline Monkiewicz, Carey Hanlin, Janee Cadlett-Jete, and Natalia Riddick for mouse colony management. We thank Michael Emanuele for guidance and suggestions on the ubiquitin remnant profiling protocol.

